# DNA damage and nuclear morphological changes in cardiac hypertrophy are mediated by SNRK through actin depolymerization

**DOI:** 10.1101/2023.07.14.549060

**Authors:** Paulina Stanczyk, Yuki Tatekoshi, Jason S. Shapiro, Krithika Nayudu, Yihan Chen, Zachary Zilber, Matthew Schipma, Adam De Jesus, Amir Mahmoodzadeh, Ashley Akrami, Hsiang-Chun Chang, Hossein Ardehali

## Abstract

**BACKGROUND:** Proper nuclear organization is critical for cardiomyocyte (CM) function, as global structural remodeling of nuclear morphology and chromatin structure underpins the development and progression of cardiovascular disease. Previous reports have implicated a role for DNA damage in cardiac hypertrophy, however, the mechanism for this process is not well delineated. AMPK family of proteins regulate metabolism and DNA damage response (DDR). Here, we examine whether a member of this family, SNF1-related kinase (SNRK), which plays a role in cardiac metabolism, is also involved in hypertrophic remodeling through changes in DDR and structural properties of the nucleus.

**METHODS:** We subjected cardiac specific (cs)-*Snrk*^-/-^ mice to trans-aortic banding (TAC) to assess the effect on cardiac function and DDR. In parallel, we modulated SNRK *in vitro* and assessed its effects on DDR and nuclear parameters. We also used phospho-proteomics to identify novel proteins that are phosphorylated by SNRK. Finally, co-immunoprecipitation (co-IP) was used to verify Destrin (DSTN) as the binding partner of SNRK that modulates its effects on the nucleus and DDR.

**RESULTS:** cs-*Snrk*^-/-^ mice display worse cardiac function and cardiac hypertrophy in response to TAC, and an increase in DDR marker pH2AX in their hearts. Additionally, *in vitro Snrk* knockdown results in increased DNA damage and chromatin compaction, along with alterations in nuclear flatness and 3D volume. Phospho-proteomic studies identified a novel SNRK target, DSTN, a member of F-actin depolymerizing factor (ADF) proteins that directly binds to and depolymerize F-actin. SNRK binds to DSTN, and DSTN downregulation reverses excess DNA damage and changes in nuclear parameters, in addition to cellular hypertrophy, with SNRK knockdown. We also demonstrate that SNRK knockdown promotes excessive actin depolymerization, measured by the increased ratio of globular (G-) actin to F-actin. Finally, Jasplakinolide, a pharmacological stabilizer of F-actin, rescues the increased DNA damage and aberrant nuclear morphology in SNRK downregulated cells.

**CONCLUSIONS:** These results indicate that SNRK is a key player in cardiac hypertrophy and DNA damage through its interaction with DSTN. This interaction fine-tunes actin polymerization to reduce DDR and maintain proper CM nuclear shape and morphology.

**Clinical Perspective:**

1. *What is new?*

- Animal hearts subjected to pressure overload display increased SNF1-related kinase (SNRK) protein expression levels and cardiomyocyte specific SNRK deletion leads to aggravated myocardial hypertrophy and heart failure.
- We have found that downregulation of SNRK impairs DSTN-mediated actin polymerization, leading to maladaptive changes in nuclear morphology, higher DNA damage response (DDR) and increased hypertrophy.
2. *What are the clinical implications?*

- Our results suggest that disruption of DDR through genetic loss of SNRK results in an exaggerated pressure overload–induced cardiomyocyte hypertrophy.
- Targeting DDR, actin polymerization or SNRK/DSTN interaction represent promising therapeutic targets in pressure overload cardiac hypertrophy.

## INTRODUCTION

Cardiac left ventricular (LV) hypertrophy (LVH) is characterized by increased thickness of the LV. Although physiological cardiac hypertrophy may occur in response to exercise, pathological hypertrophy commonly occurs in response to pressure overload and is associated with diseases such as hypertension, valvular disease, and ischemic heart disease (1). In many of these cases, the LVH eventually leads to heart failure. Over the past several decades, several proteins, processes and pathways have been identified to mediate cardiac hypertrophy. Recently, Sadek’s group identified that DNA damage response (DDR) plays a major role in the development of cardiac hypertrophic growth (2), however, the mechanism of how DDR occurs in cardiac hypertrophy is not known.

During states in which energy is depleted, shift of AMP/ADP to ATP ratio cause activation of AMPK in the heart to increase energy production from lipid oxidation, glycolysis, and glycogen breakdown (3). In addition to lipid and glucose catabolism, AMPK also increases mitochondrial biogenesis to enhance the production of ATP (4). SNRK is a serine-threonine kinase and a mammalian protein relative of AMPK. SNRK contains an ATP-binding domain and a serine/threonine kinase domain with a conserved T-loop residue at its N-terminal end, highly homologous to AMPK kinase domain (5). However, unlike AMPK, SNRK does not require an additional stimulus for activation such as increased AMP/ATP ratio (6). Expression of SNRK mRNA is widespread in the tissues of adult mice, rats, and humans, as indicated by Northern blotting and *in situ* analysis (5, 7, 8). Since SNRK has sequence similarity to AMPK, its cellular role likely resembles that of other proteins in the AMPK family, with SNRK signaling partners and pathways emerging as new areas of research. To date, it has been suggested that SNRK may be involved in neuronal apoptosis (7), blood vessel development (9–11), and kidney inflammation (12). SNRK has also been shown to play a role in normal physiology and disease in highly metabolic tissues such as adipose (13–15) and cardiac (16, 17) tissues. We have also shown that SNRK inhibits proliferation of colon cancer cells (18) and improves cardiac mitochondrial efficiency and decreases mitochondrial uncoupling in the heart (19).

While AMPK is an established metabolic regulator, it functions also as a sensor of genomic stress (20, 21). AMPK is confined and activated in the cytosol in response to low nutrients (22, 23), while oxidant exposure facilitates the AMPK nuclear translocation (23, 24). Importantly, nuclear AMPK has been involved in re-organization of nucleoli, allowing AMPK to modulate cell proliferation and apoptosis (25). Interestingly, LKB1 (AMPK and SNRK activator (26)), can also shuttle between cytoplasm and nucleus (27), indicating a multifaceted mechanism to control AMPK family localization and activation, and that SNRK might also participate in the nuclear processes. SNRK was originally found to be upregulated and localized to the nucleus in response to apoptosis in rat cerebellar granule neurons (7). Further, SNRK suppresses adipocyte inflammation, and SNRK may phosphorylate histone deacetylase 1 (14, 28), a known epigenetic regulator of cellular proliferation and apoptosis (29).

Members of the cofilin family of actin depolymerizing factor proteins regulate actin dynamics through depolymerizing actin filaments (30). There are three highly conserved cofilin proteins: Cofilin-1 (CFL1; also known as non-muscle-or n-Cofilin), Cofilin-2 (CFL2; also known as muscle-or m-Cofilin), and Destrin (DSTN; also known as actin-depolymerizing factor (ADF)). Although these proteins have distinct expression patterns, they share the function of binding to globular and filamentous actin (G- and F-actin, respectively) (31), resulting in an increase in the number of actin monomers and filament fragments (32). In addition to regulating cell shape, motility, and sarcomeric function in myocytes, cytosolic actin can regulate nuclear size, shape, and polarity through interactions with tethering complexes embedded in the nuclear envelope (33). These tethering complexes known as linker of nucleoskeleton and cytoskeleton (LINC) have been shown to play pivotal roles in cellular mechano-transduction and DDR (34, 35). Similarly, knockdown of cofilin-family proteins results in enlarged and abnormally shaped nuclei and increased DNA damage (36).

In this paper, we show that SNRK plays a major role in hypertrophy mediated by DDR. We also show that this effect is mediated through a member of the cofilin family of proteins, DSTN. SNRK directly binds to DSTN and this interaction leads to changes in actin polymerization, which affects both nuclear morphology and DDR response.

## METHODS

### Animal studies

All studies adhered to the National Institutes of Health Guide for the Care and Use of Laboratory Animals; the protocols were approved by Institutional Animal Care and Use Committee of Northwestern University.

### Phospho-proteomic studies

Phospho-proteomic data are available in the PRIDE repository at https://www.ebi.ac.uk/pride/.

### Statistical analysis

Data are presented as mean ± standard error mean (SEM). Statistical significance was assessed with Student’s t-test and ANOVA with post hoc Tukey’s or Holm-Sidak’s test for multiple group comparison, as indicated in the Figure legends. All analysis were conducted using GraphPad Prism v9. A P-value less than 0.05 was considered statistically significant.

## RESULTS

### Deletion of Snrk leads to cardiac hypertrophy

We first assessed whether SNRK levels increase in cardiac hypertrophy. We first measured the levels of SNRK protein in the heart 4 weeks after pressure overload due to TAC and observed higher SNRK protein levels (**Figure 1A and 1B**). To assess the role of SNRK in cardiac hypertrophy, we measured cell size after *Snrk* knockdown (KD) in H9c2 cells and cardiac size and function in cardiac-specific *Snrk* knockout (KO) mice (cs-*Snrk*^-/-^) subjected to TAC. We first showed that *Snrk* KD in H9c2 cells results in an increase in cell size (**Figure S1**). Although there was no difference in cardiac size at baseline, cs-*Snrk*^-/-^ mice displayed larger hearts and higher heart weight/tibial length (HW/TL) after TAC compared to littermate controls (**Figure 1C and 1D**). Exaggerated cardiac hypertrophy after TAC in the hearts of cs-*Snrk*^-/-^ mice was demonstrated by echocardiography, as these mice displayed higher interventricular septal thickness end-diastole (IVSd) (**Figure 1E**) and posterior LV wall thickness end-diastole (PWTd) (**Figure 1F**). cs-*Snrk*^-/-^ mice also displayed a significantly lower ejection fraction (EF) and fractional shortening (FS) 3-weeks after TAC (**Figure 1G and 1H**). There was no difference in cardiac function and other cardiac parameters between male and female wild type (WT) or cs-*Snrk*^-/-^ mice after sham and TAC surgery (**Figure S2**). Additionally, we noted no difference in cardiac function (as assessed by echocardiography) and heart rate between WT and αMHC-Cre (used for tissue-specific deletion of *Snrk* in the heart) after sham and TAC surgery (**Figure S3**). These results indicate that SNRK protein levels are increased in TAC-induced cardiac hypertrophy and its deletion leads to an increase in cardiac hypertrophy and eventually cardiac failure in response to pressure overload.

**Figure 1.**
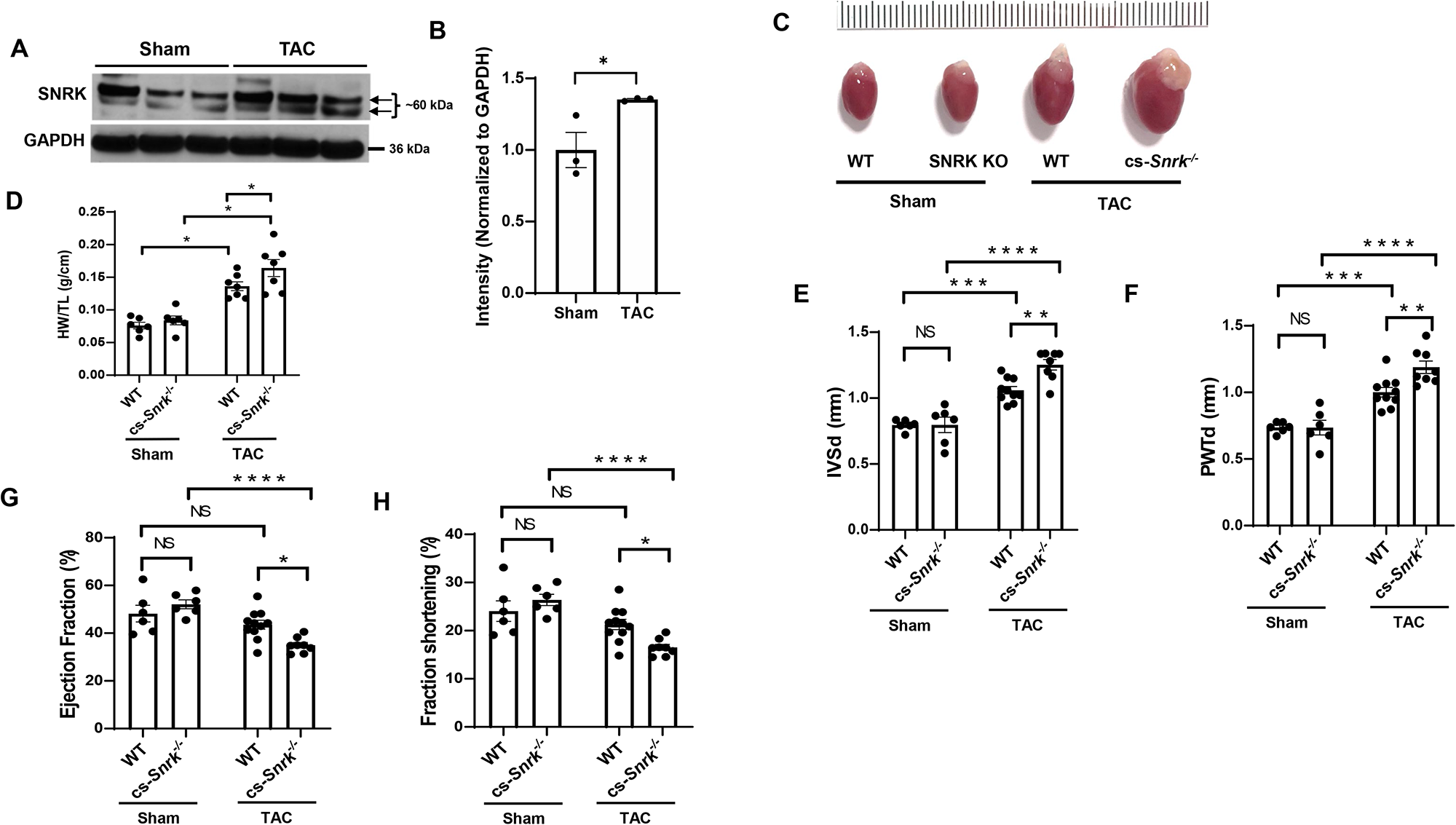
SNRK is increased in cardiac hypertrophy and its deletion leads to exaggerated hypertrophy. **A,** SNRK protein levels in heart tissue from mice subjected to sham or 4-weeks after TAC operation. **B,** Summary bar graph summary of Panel A (n=3 animals per condition, unpaired t-test, *P<0.05). **C**, Image of freshly extracted hearts from WT and cs-*Snrk*^-/-^ mice 4-weeks after sham operation or TAC. Scale bar = 0.1 cm. **D**, HW/TL in WT and cs-*Snrk*^-/-^ mice 4-weeks after sham operation or TAC (n>6 animals per condition, two-way ANOVA with Tukey’s test, *P<0.05). **E-F**, IVSd (E) and PWTd (F) in WT and cs-*Snrk*^-/-^ mice 4-weeks after sham operation or TAC (n=6-10, two-way ANOVA with Tukey’s multiple comparisons test). **G-H**, EF (G) and FS (H) in WT and cs-*Snrk*^-/-^ mice 3-weeks after sham operation or TAC (n=3-11, two-way ANOVA with Sidak’s multiple comparisons test, *P<0.05, **P<0.01, ***P<0.001, ****P<0.0001.

### SNRK mediates DDR and alters nuclear morphology

Since SNRK levels are increased in cardiac hypertrophy and cs-*Snrk*^-/-^ mice show higher cardiac hypertrophy in pressure overload, and DDR has been implicated in cardiac hypertrophy (2), we then asked whether SNRK plays a role in DDR response. We first treated H9c2 cells with *Snrk* siRNA and assessed pH2AX protein and 8-Hydroxyguanosine (8-OHdG) levels. pH2AX was higher in cells with *Snrk* KD (**Figure 2A and 2B**) indicating increased double strand DNA break (DSB). Additionally, 8-OHdG was increased with *Snrk* siRNA, indicating increased DNA damage (**Figure 2C and 2D**). We also assessed DSB and DDR by measuring pH2AX and phosphorylated ATR (p-ATR) in the hearts of cs-*Snrk*^-/-^ mice with sham operation or after TAC, and noted a significant increase in pH2AX levels (**Figure 2E and 2F**) and p-ATR/total ATR (**Figure 2G and 2H**) in the hearts of cs-*Snrk*^-/-^ mice at baseline and after TAC.

**Figure 2.**
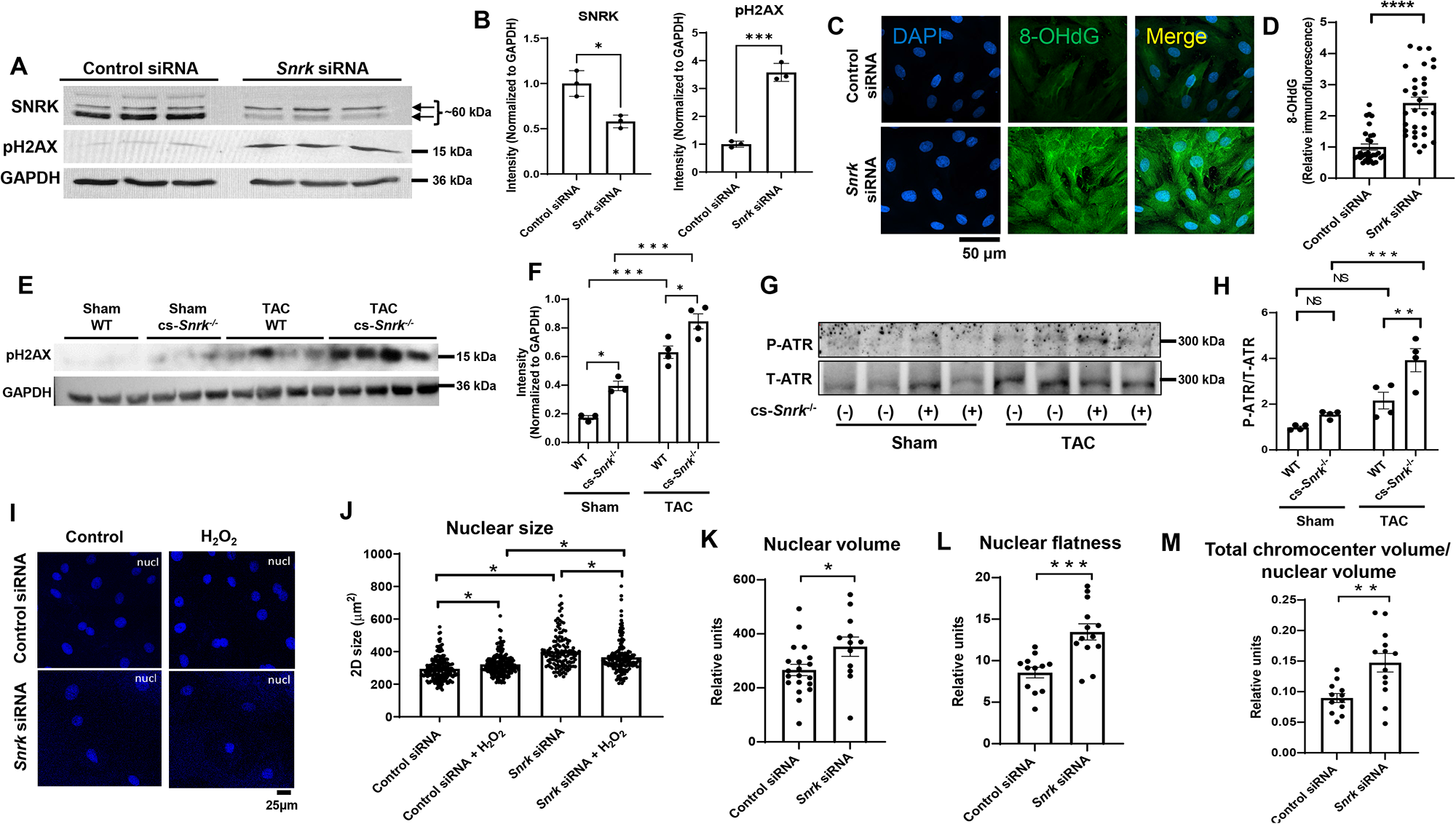
SNRK modulates DDR and nuclear morphology. **A,** Western blot of SNRK and pH2AX in H9c2 cells treated with control or *Snrk* siRNA. **B,** Summary bar graph of Panel A (n=3, Student’s t-test). **C,** Representative IF images of 8-OHdG stain of H9c2 cells treated with control or *Snrk* siRNA. Scale bar = 50 µm. **D,** Results of the 8-OHdG fluorescence from Panel C (n≥30 cells measured for each condition, Student’s t-test). **E,** Western blot of pH2AX in the hearts of WT and cs-*Snrk^-/-^* mice treated with sham or after TAC. **F,** Summary bar graph of the results in Panel E (n=3-4, two-way ANOVA with Tukey’s test). **G,** Western blot of p-ATR and total ATR in H9c2 cells treated with control or *Snrk* siRNA. **H,** Summary bar graph of Panel G (n=4, two-way ANOVA with Tukey’s test). **I,** Representative IF images of DAPI stain of H9c2 cells treated with control or *Snrk* siRNA. Scale bar = 25 µM. **J,** Results of the nucleus size measurements from Panel I (n=3 independent experiments, with ≥42 cells measured for each condition per replicate, Student’s t-test). **K-M,** Nuclear volume (K), nuclear flatness (L), and total chromocenter volume/nuclear volume (M) measurements in H9c2 cells treated with control or *Snrk* siRNA. Nuclear flatness reflects the ratio of the length of the intermediate axis of the cell to length of the shortest axis. (n ≥12 cells measured for each condition, Student’s t-test). *P<0.05, **P<0.01, ***P<0.001, ****P<0.0001.

Increased DNA damage has been associated with alteration in nuclear morphology and subnuclear organization (37, 38), so we also assessed the effects of SNRK on nuclear size and noted that deletion of *Snrk* in H9c2 cells results in a significant increase in 2D nuclear size (**Figure 2I and 2J**). We then measured nuclear morphology by performing 3D nuclear parameter analysis and measuring nuclear volume and number of chromocenters in H9c2 cells with *Snrk* KD. It should be noted that one of the consequences of nuclear DNA damage is actin-mediated heterochromatin restructuring (37, 38), which is the reason we included this parameter in our analysis. We observed that *Snrk* downregulation significantly increased nuclear volume (**Figure 2K**), nuclear flatness (**Figure 2L**), and total chromocenter volume / nuclear volume (**Figure 2M**). These results suggests that *Snrk* KD results in nuclear instability, which could potentially be through increased DNA damage and heterochromatin restructuring.

### SNRK interacts with DSTN

To determine the mechanism by which SNRK mediates its effect on nuclear structure and DDR, we performed phospho-proteomic analysis with overexpression of either an empty vector (EV), mutant D158A kinase-dead SNRK construct (SNRK-Mut), or WT SNRK. Two independent phospho-proteomic datasets were prepared, each containing phospho-peptides corresponding to cells treated with EV, SNRK-Mut, and WT (**Figure S4A**). Database 1 yielded 1400 unique phospho-peptides in total, filtered down to only the phosphorylated sites that can be assigned to a single amino acid with >75% in at least 1 sample. That yielded 656 phospho-peptides quantified on 435 proteins. For database 2, 2656 unique phospho-peptides in total were identified, and after filtering, it yielded 1480 phospho-peptides quantified on 772 proteins. A significant upregulation in phosphorylation of several proteins was found in cells overexpressing WT SNRK compared to EV or SNRK-Mut (**Figure 3A, S4B**), with majority corresponding to nuclear processes based on gene ontology (GO) analysis (**Figure 3B**) and STRING analysis (**Figure S4C**). Several proteins were significantly enriched among proteins that displayed increased phosphorylation in cells overexpressing WT SNRK compared to EV or WT vs SNRK-Mut (**Figure 3A, S4B**), with differences in DSTN S3 phosphorylation in WT vs both EV and SNRK-Mut groups (**Figure 3C**). Because of this, and the fact that DSTN has a well-established role in actin remodeling (30–32), and that actin is an important mediator of nuclear structure and DDR (34), we selected this protein for further study.

**Figure 3.**
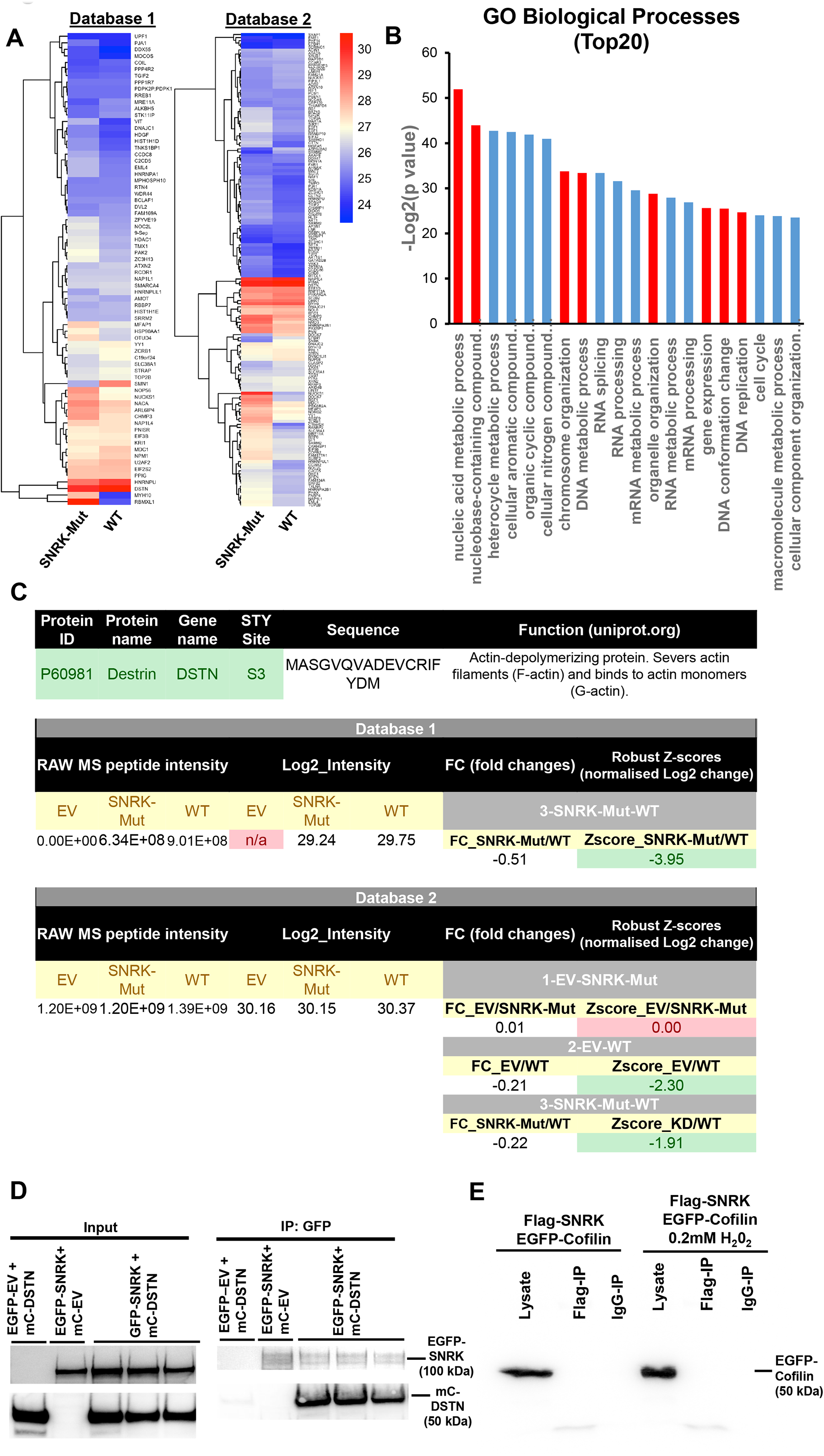
SNRK interacts with DSTN. **A,** Heatmap summary of all identified peptides and their corresponding log2 intensity values from the phosphoproteomic study, with the values above 1.5 and below 1.5 z-scores (normalized Log2 fold changes) for Mutant vs WT for both databases 1 and 2. **B,** Summary of Gene ontology (GO) analysis of top 20 pathways of all upregulated and downregulated genes identified as upregulated in WT vs Mutant and WT vs EV from both databases, performed using http://www.pantherdb.org/. Red bars represent [rocesses related to cell cycle, and organelle and chromosome organization. **C,** Summary of DSTN S3 raw phosphoproteomic data and corresponding log2 intensity and Z-score fold changes from both databases. **D,** Co-IP studies of overexpressed constructs with EGFP fusion to SNRK and DSTN fused to m-Cherry probed with GFP and mCherry antibodies. The co-IP studies demonstrated that DSTN binds to SNRK. **E,** Co-IP studies of overexpressed constructs with FLAG fusion with SNRK and cofilin-1 fused to GFP (probed with GFP antibody). IP experiments were done with FLAG antibody.

We then performed co-immunoprecipitation (co-IP) studies to determine whether SNRK and DSTN can directly interact and/or form a protein complex. For these studies, we overexpressed constructs with EGFP fusion to SNRK (EGFP-SNRK) and m-cherry fused to DSTN (mc-DSTN), as well as empty vector constructs (EGFP-EV and mc-EV, respectively). The co-IP studies demonstrated that while both SNRK/DSTN and reverse DSTN/SNRK pull-downs can be detected, there was no signal for EGFP/DSTN nor mCherry/SNRK combination (**Figure 3D, and Figure S5A**), indicating that SNRK and DSTN physically interact with each other. Since DSTN belongs to cofilin family of proteins (30), we also tested if SNRK binds to cofilin-1 as well. Our results indicated that there is no interaction between SNRK and cofilin-1 (**Figure 3E and Figure S5B**), indicating that SNRK only binds to DSTN and not to cofilin-1.

### DSTN mediates the effects of SNRK on DDR and nuclear morphology

Since SNRK binds to DSTN, we next studied whether the effects of SNRK on nuclear morphology and DDR are mediated through DSTN. Thus, we downregulated *Dstn* in the presence of *Snrk* KD to study the role of DSTN in SNRK-mediated changes in DDR and nuclear morphology. We first showed that *Dstn* KD reverses the effects of *Snrk* KD on cell hypertrophy by reducing cell size (**Figure 4A and 4B**). We then tested whether DSTN has an effect on DNA damage (by measuring 8-OHdG) and DSB (by measuring pH2AX) and demonstrated that *Dstn* KD reverses the increase in 8-OHdG and pH2AX signal as a result of *Snrk* KD (**Figure 4C-F**). We also assessed DDR by measuring pATM and showed that a reduction in DSTN also reverses the markers of DDR induced by *Snrk* KD (**Figure 4G and 4H**). Additionally, *Dstn* KD reversed the effects of *Snrk* KD on 3D nuclear volume and nuclear flatness (**Figure 4I-K**). These results indicate that the effects of *Snrk* KD on DNA damage, DDR, nuclear volume and flatness are mediated through DSTN.

**Figure 4.**
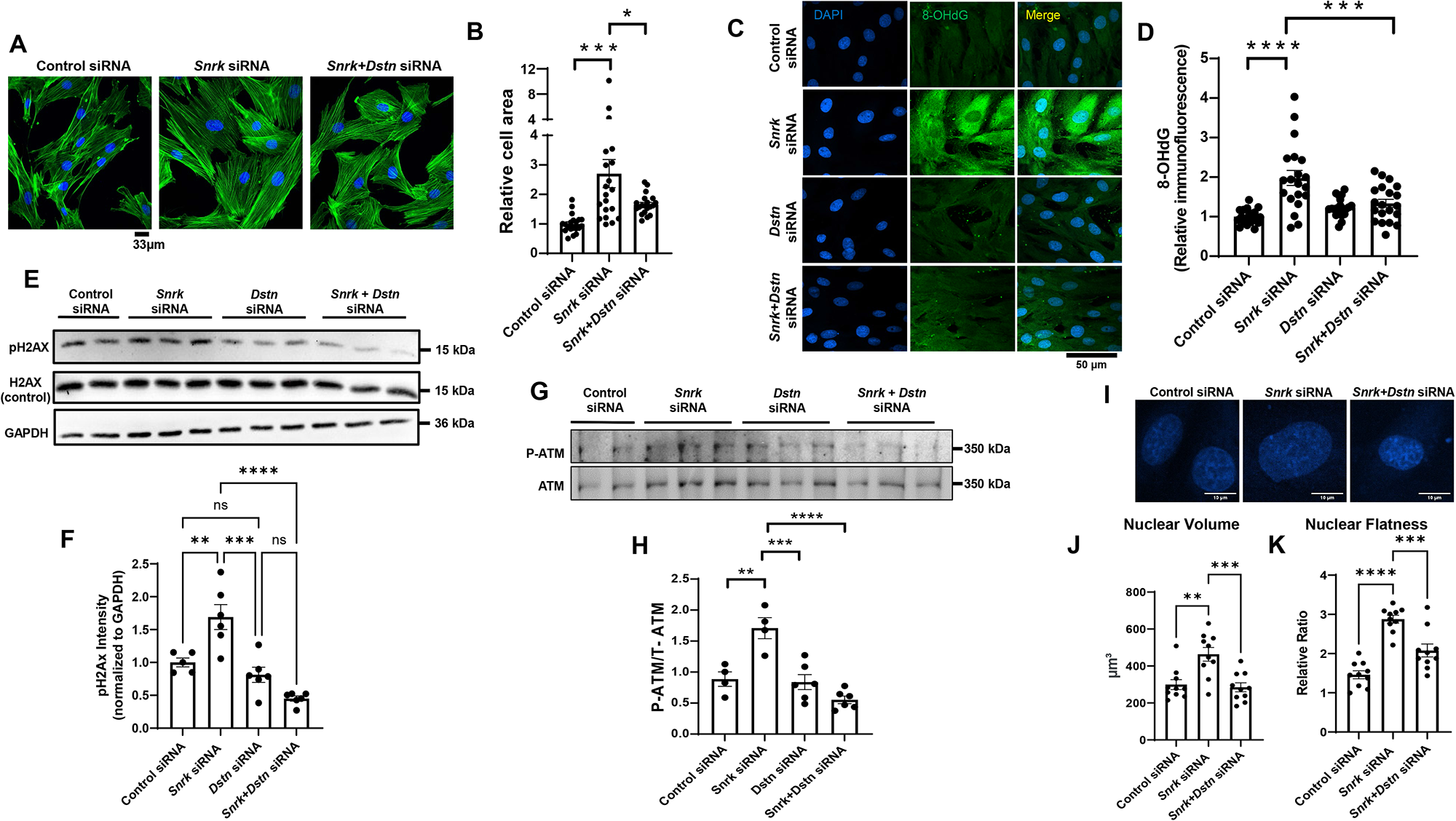
DSTN KD reverses the DDR and changes in nuclear morphology induced by SNRK KD. **A**, Representative IF images of H9c2 cells treated with control or *Snrk* siRNA alone, or combination of *Snrk* and *Dstn* siRNA and stained with Phalloidin-iFluor 488 to assess cell surface area. **B,** Summary of results in Panel A (n=20 cells for each condition, one-way ANOVA with Tukey’s test). **C,** Representative IF images of 8-OHdG stain of H9c2 cells treated with control, *Snrk*, *Dstn*, or *Snrk+Dstn* siRNA. Scale bar = 50 µM. **D,** Results of the 8-OHdG fluorescence from Panel C (n=20 cells measured for each condition, two-way ANOVA with Tukey’s test). **E,** Western blot of SNRK, DSTN, pH2AX, and H2AX in H9c2 cells treated with control siRNA, *Snrk* siRNA, *Dstn* siRNA, or *Snrk*+*Dstn* siRNA. **F,** Summary of results in Panel E (n=3, one-way ANOVA with Tukey’s test). **G,** Western blot of p-ATM and total ATM in H9c2 cells treated with control, *Snrk*, *Dstn*, or *Snrk+Dstn* siRNA. **H,** Summary bar graph of Panel G (n=4-6, two-way ANOVA with Tukey’s test). **I,** Representative IF images of H9c2 cells treated with DAPI staining and with control, *Snrk*, or *Snrk*+*Dstn* siRNA. Scale bar = 10 µm. **J-K,** Nuclear volume (J) and nuclear flatness (K) in H9c2 cells treated with control, *Snrk*, or *Snrk*+*Dstn* siRNA. Nuclear flatness reflects the ratio of the length of the intermediate axis of the cell to length of the shortest axis. (n=10, one-way ANOVA with Tukey’s multiple comparisons test). *P<0.05, **P<0.01, ***P<0.001, ****P<0.0001.

### SNRK affects DDR through F-actin depolymerization mediated by DSTN

DSTN and cofilin proteins mediate F-actin depolymerization and proper nuclear tethering to the actin cytoskeleton for maintenance of nuclear morphology (34). Thus, we next studied whether SNRK alters F-actin depolymerization through DSTN. An increase in the G-vs F-form of actin is associated with increased actin depolymerization and changes in nuclear morphology (36). We showed that *Snrk* KD is associated with an increase in G/F actin ratio, which was reversed with *Dstn* KD (**Figure 5A and 5B**). Additionally, we measured the alignment of the nuclear long axis with the direction of cytosolic actin filaments, which is used as a marker of appropriate nuclear-cytoskeletal tethering (33, 39). Our results showed that treatment of H9c2 cells with *Snrk* siRNA disrupted nuclear axis to F-actin angle (**Figure 5C and 5D**), which was reversed with *Dstn* KD (**Figure 5C and 5D**). Additionally, phalloidin fluorescence, which specifically binds to F-actin (40), was reduced with *Snrk* siRNA, and this was also reversed with *Dstn* KD (**Figure 5E**). Finally, to demonstrate the effect of *Snrk* KD was through changes in actin depolymerization, we treated *Snrk* KD cells with the F-actin stabilizing drug Jasplakinolide (JAS) (41). Initial dose curve experiments demonstrated that 30nM JAS was sufficient to increase G/F actin ratio without independently causing DNA damage H9C2 cells (**Figure S6**). We also demonstrated that while *Snrk* KD reduces F-actin fluorescence, JAS reverses this effect (**Figure 5F and 5G**), confirming that JAS increased actin polymerization. Similar to the results with *Dstn* KD, JAS treatment reversed the increase in pH2AX and pATM levels with *Snrk* KD (**Figure 5H-K**), indicating that the increase in DNA damage by *Snrk* KD is through actin polymerization. Finally, treatment of cells with JAS also reversed the increase in nuclear volume and flatness caused by *Snrk* KD (**Figure 5L and 5M**).

**Figure 5.**
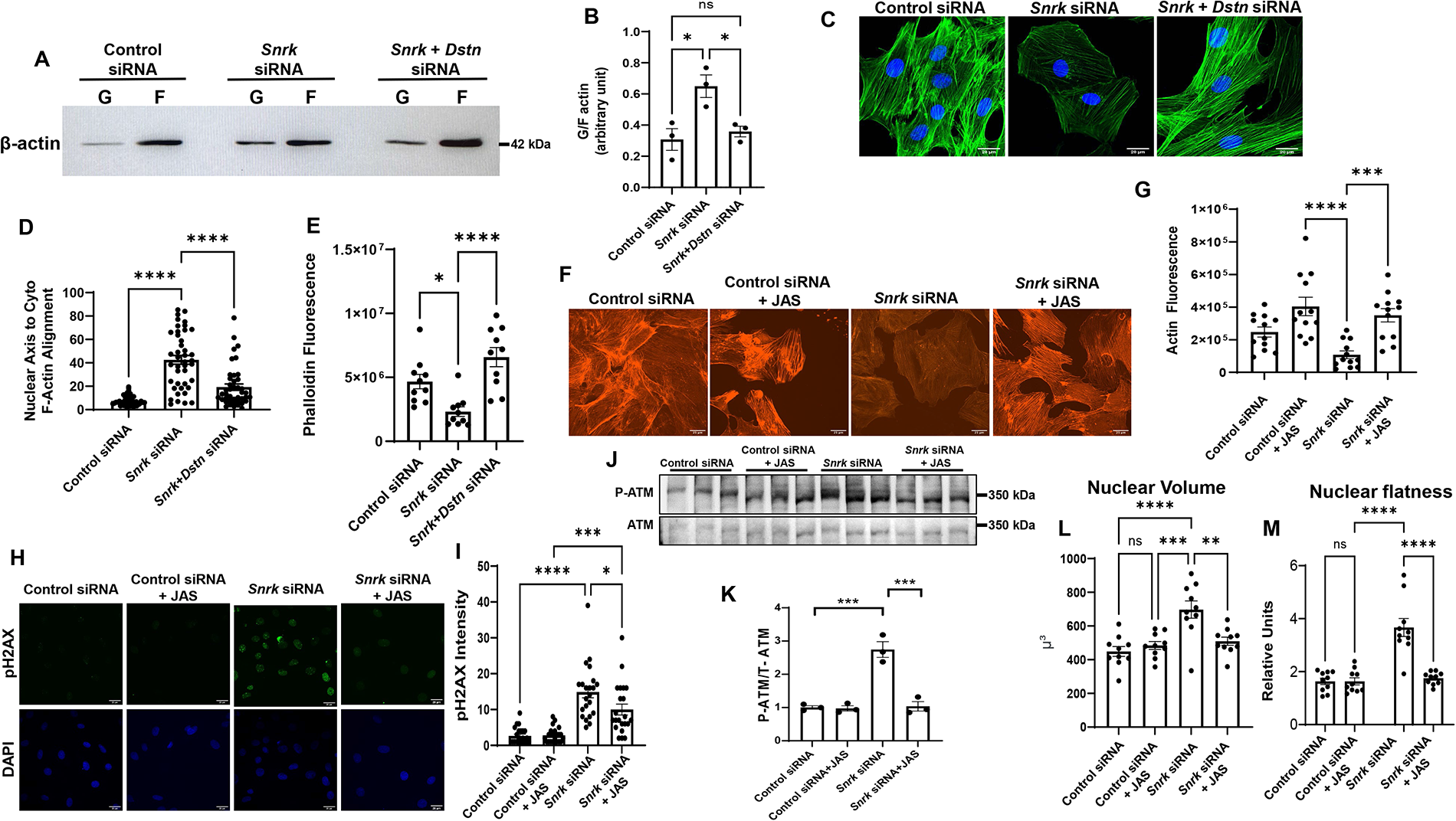
SNRK alters DDR and nuclear morphology through actin depolymerization. **A,** Representative Western blot of F and G actin in H9c2 cells treated with control, *Snrk* or *Snrk*+*Dstn* siRNA. **B,** Summary bar graph of Western blot results in A (n=3, one-way ANOVA with Tukey’s test). **C,** Representative IF images probing with actin staining in H9c2 cells treated with control, *Snrk* or *Snrk*+*Dstn* siRNA. Scale bar = 20 µm. **D,** Summary bar graph of nuclear-actin angle in H9c2 cells treated with control, *Snrk* or *Snrk*+*Dstn* siRNA (n=42, one-way ANOVA with Tukey’s Multiple Comparisons Test). **E,** Phalloidin fluorescence in H9c2 cells treated with control, *Snrk* or *Snrk*+*Dstn* siRNA (n=10, one-way ANOVA with Tukey’s test). **F,** Representative fluorescence actin images of H9c2 cells treated with *Snrk* siRNA in the presence and absence of JAS. Scale bar = 20 µm. **G,** Summary bar graph of actin fluorescence of H9c2 cells with Snrk siRNA and JAS (n=12, one-way ANOVA with Tukey’s Multiple Comparisons Test). **H,** Representative pH2AX immunohistochemistry of H9c2 cells treated with *Snrk* siRNA in the presence and absence of JAS. Scale bar = 25 µm. **I,** Summary bar graph of pH2AX staining of H9c2 cells with *Snrk* siRNA and JAS (n=22, one-way ANOVA with Tukey’s Multiple Comparisons Test). **J,** Western blot of p-ATM and total ATM in H9c2 cells treated with control or *Snrk* siRNA in the presence and absence of JAS. **K,** Summary bar graph of Panel G (n=3, two-way ANOVA with Tukey’s test). **L-M,** Nuclear volume (L) and nuclear flatness (M) in H9c2 cells treated with *Snrk* siRNA in the presence and absence of JAS. Nuclear flatness reflects the ratio of the length of the intermediate axis of the cell to length of the shortest axis. (n=10, one-way ANOVA with Tukey’s test). *P<0.05, **P<0.01, ***P<0.001, ****P<0.0001.

We next conducted *in vivo* experiments to confirm the role of DDR in the exaggerated cardiac hypertrophy with *Snrk* deletion by treating WT and cs-*Snrk*^-/-^ mice with an ATM inhibitor, KU-60019, and assessing cardiac function. cs-*Snrk^-/-^* mice displayed lower EF and FS and larger PWTd and LV diameter end-systolic (LVDs) than WT after TAC, but the changes were mitigated by the treatment of KU-60019 (**Figure 6A-D**). Collectively, these data indicate that SNRK regulates nuclear morphology and DDR through DSTN-mediated actin depolymerization, and that deletion of *Snrk* leads to cardiac hypertrophy through alterations of this pathway (**Figure 6E**).

**Figure 6.**
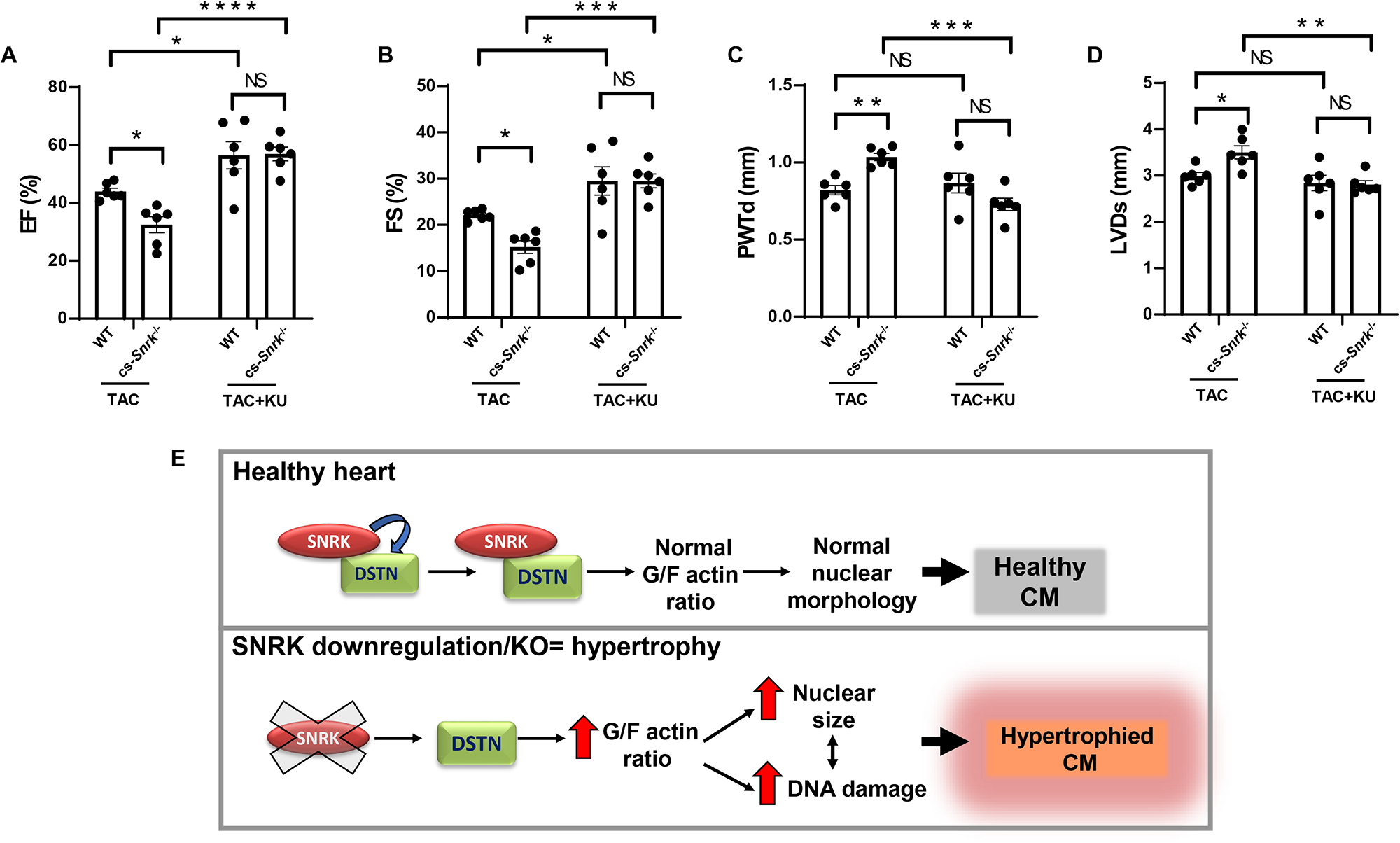
Inhibition of DDR signaling inhibits pressure overload–induced cardiomyocyte hypertrophy in cs-*Snrk*^-/-^. **A-D**, EF (A), FS (B), PWTd (C), and LVDs (D) in WT and cs-*Snrk*^-/-^ mice 4-weeks after sham operation or TAC with or without KU-60019 (n=6, two-way ANOVA with Holm-Sidak’s multiple comparisons test). *P<0.05, **P<0.01, ***P<0.001, ****P<0.0001. **E,** Schematic model for the SNRK-DSTN interaction in relation to SNRK KO hypertrophy. SNRK regulates nuclear morphology and DDR through DSTN mediated actin depolymerization, and that deletion of *Snrk* leads to cardiac hypertrophy through perturbation of G/F actin ratio.

## DISCUSSION

Pathological cardiac hypertrophy can lead to diastolic dysfunction and subsequently to heart failure (1). Recently, Nakada *et al.* showed that DDR plays a major role in the progression of cardiac hypertrophy, but the exact mechanism by which DDR is induced is not clear (2). In this paper, we studied the role of a member of the AMPK family, SNRK, in DDR and hypertrophy. We showed that *Snrk* KO mice are more susceptible to cardiac hypertrophy, and that changes in SNRK levels can modify pH2AX marker of DDR and nuclear morphology. This regulation is through SNRK interaction with DSTN, which regulates F-actin depolymerization. Thus, our data indicates that SNRK regulates cardiomyocyte homeostasis and hypertrophy in part through changes in actin polymerization status and its subsequent effects on the nucleus, as summarized in **Figure 6E**.

DDR mediates a number of pathological and physiological conditions. Shortly after birth, murine heart can recover its function after injury due to proliferation of preexisting cardiomyocytes, which is mediated through DDR (42, 43). Additionally, cardiac hypertrophy is associated with an increase in double strand breaks (DSB), and phosphorylated ataxia telangiectasia (ATM), and inhibitors of ATM kinase or deletion of ATM in cardiomyocytes reversed cardiac hypertrophy in response to TAC or angiotensin infusion (2). Similar results were obtained by Higo et al. (44), however, this group only observed single-strand DNA breaks, not DSB. Nevertheless, these studies raise the possibility of reversing cardiac hypertrophy by targeting DDR. Understanding the pathway that leads to DDR in cardiac hypertrophy can provide additional potential therapeutic targets. Our results suggest that targeting SNRK activity or its interaction with DSTN, in addition to actin polymerization may be used to reverse the process of pathological hypertrophy.

The interplay between chromatin structure, factors that regulate chromatin organization and cytoskeletal forces are essential contributors to the physical properties of the nucleus (34), and previous studies have shown alterations in cardiomyocyte nuclear morphology and size in response to stress. Cardiomyocytes display a shift of chromatin distribution to low-density with concomitant increase in nuclear size in response to stress (45). Additionally, morphological nuclear defects such as irregular nuclear envelope, enlarged and misshaped nuclei, indentations, and irregularly distributed chromatin are described in histological assessment of samples from patients with cardiomyopathy (46, 47). Changes in actin polymerization and LINC around the nucleus also reduces nuclear volume and chromatin accessibility, resulting in reduced reprogramming of pluripotent stem cells (48). Further, the importance of proper nuclear architecture is evinced by range of cardiac pathologies associated with various mutations in LMNA gene, key components in providing structural support to the nucleus, where even patients with the same mutation can exhibit a variety of heart disease symptoms, including DCM, arrhythmias and sudden cardiac death (49, 50). We demonstrate a novel role for SNRK-DSTN interaction on DDR and nuclear morphology through regulation of actin filament turnover, providing a novel mechanism for DDR and nuclear structure in the heart.

Although we provide a novel mechanism, there are some limitations with our study. First, we did not identify transcriptional alterations that occur due to changes in nuclear morphology and DDR that may mediate the protective effects of SNRK. Additionally, given the role of SNRK in cardiac metabolism and cardioprotection (16, 17, 51), it is possible that there is a functional link between metabolic and DDR changes that occur in response to SNRK in the heart. Finally, although we have *in vitro* data demonstrating that alterations in global actin depolymerization rescues the effects of *Snrk* KD on nuclear morphology, we did not alter actin depolymerization in the heart of cs-*Snrk*^-/-^ mice to show such intervention reverses cardiac hypertrophy.

In summary, our results demonstrate that the DDR associated with cardiac hypertrophy is mediated by SNRK. This effect is through SNRK interaction with a member of the ADF family of proteins, DSTN. SNRK interaction with DSTN leads to changes in actin polymerization, which affect nuclear morphology and DDR response. These studies suggest that targeting DDR, actin polymerization or SNRK/DSTN interaction may have therapeutic potentials in pressure overload cardiac hypertrophy.

## Acknowledgements

We would like to thank Meng Shang, Eric Xia, Tivoli Nguyen and Mingyang Liu for managing our mouse colony and genotyping the mice used in this study, Chunlei Chen for conducting the animal surgery experiments, and Zeinab Najafi for her technical assistance. We thank Young Ah Goo from the Northwestern University Proteomics Center of Excellence for running the mass-spectrometry and MaxQuant analysis for our proteomics experiments. We also acknowledge Dr. Jin Jing and Dr. Pan Liu for their help with design and analysis of phosphoproteomics studies.

## SUPPLEMENTARY FIGURE LEGENDS

**Figure S1. SNRK knockdown results in cell hypertrophy in H9c2 cells.** Representative live bight field (BF) and fluorescent images and quantification of H9c2 cells treated with control or *Snrk* siRNA alone, stained with cell tracker CM-DiI (red/cell size staining) and Hoechst 33342 (blue/nuclear stain). n>30 cells measured for each condition, one-way ANOVA with Tukey’s test, *P<0.05.

**Figure S2. Cardiac function of male and female cs-*Snrk*^-/-^ mice after Sham and TAC surgery. A-F,** HR (A), EF (B), FS (C), LVDs (D), LVDd (E), IVSd (F), and PWTd (G) in cs-*Snrk*^-/-^ mice after sham and TAC surgery (n≥3)

**Figure S3. Cardiac function of WT and αMHC-cre mice after Sham and TAC surgery. A-G,** HR (A), EF (B), FS (C), LVDs (D), LVDd (E), IVSd (F), and PWTd (G) in WT and αMHC cre mice after sham and TAC surgery. (n≥5)

**Figure S4. Phosphoproteomic studies with SNRK overexpression. A,** Venn diagram summary of phosphoproteomic analysis of overexpression model HepG2 with 48h transient overexpression of EGFP-EV, EGFP-SNRK-WT and EGFP-SNRK-Mutant from n=2 independent experiments (database 1 and database 2). **B,** Heatmap summary of all identified peptides and their corresponding log2 intensity from phosphoproteomic study) with the values above 1.5 and below 1.5 z-scores (normalized Log2 fold changes) for EV vs Mutant, EV vs WT and EV vs Mutant for both databases 1 and 2. **C,** STRING interaction network (functional protein-protein association bioinformatics) analysis of joint database 1 and 2 upregulated proteins (upregulated in WT vs Mutant and WT vs EV).

**Figure S5. Western Blots of lysates from SNRK and DSTN overexpression experiments and control co-IP experiment for SNRK-cofilin1 interaction. A,** Western blot of lysates from overexpression of constructs with EGFP fusion to SNRK and DSTN fused to m-Cherry treated at baseline and after treatment with 0.2mM H_2_O_2_. **B,** Co-IP control studies for the SNRK-cofilin1 interactions. The lysates were IPed with FLAG antibody and the membranes were also probed with FLAG antibody.

**Figure S6. JAS dose response and actin F/G actin ratio and DNA damage response. A,** Western blot of H9c2 cell extracts treated with different concentrations of JAS (actin polymerization inhibitor drug) and probed to assess the levels of F- and G-actin. **B,** Western blot of H9c2 cell extracts treated with different concentrations of JAS (actin polymerization inhibitor drug) and probed with pH2AX.

**Table S1. Database 1 raw phosphoproteomic analysis results.**

**Table S2. Database 2 raw phosphoproteomic analysis results.**

**Table S3. Upregulated proteins used for GO analysis.**

